# The marine goby – alpheid shrimp symbiosis does *not* correlate with larger fish eye size

**DOI:** 10.1101/329094

**Authors:** Klaus M. Stiefel, Rodolfo B. Reyes

## Abstract

**The symbiosis between marine gobies and Alpheid shrimp is based on an exchange of sensory performance (look-out for predators) by the goby versus muscular performance (burrow digging) by the shrimp. Using a comparative approach, we estimate the excess investment by the goby into its visual system as a consequence of the symbiosis. When correlating eye size with fish length for both shrimp-associated and solitary gobies, we find that the shrimp-associated gobies do *not* have larger eyes than size-matched solitary gobies.**

**We do find a trend, however, in that the shrimp-associated gobies live at shallower depths than the solitary gobies, indicative of the visual nature of the symbiosis. We discuss the implications of symbiosis based on large and small energy investments, and the evolutionary modifications likely necessary to include shrimp-goby communication into the behavior of the goby.**

## Introduction

Mutualistic symbiosis is an interaction between individuals of different species to the mutual benefit of both. In the tropical and sub-tropical Indo-Pacific, a mutualistic symbiosis between small benthic fishes, gobies, and alpheid shrimp exists (Fig. 1, Karplus, 1981; Karplus et al, 1981). In this symbiosis, the shrimp digs a burrow in sandy areas. A majority of the time the shrimp spends outside of the burrow it contacts the goby’s body with its antenna (Fig. 1, Moehring, 1972; Karplus, 1979; Preston, 1978). These burrows are 10s of cm long, with side tunnels and chambers, and often several entrances (Karplus et al., 1974). The shrimp, usually a pair of them, are safe in the burrows but due to poor vision (Jaafar & Zeng, 2012) would be in danger during foraging trips over the open sand. The fish communicates the presence of dangers to the shrimp via tactile cues. The cues include a tail-wiggle for minor alerts and a rushed escape into the burrow for major alerts. The symbiosis allows the pair to colonize sandy regions otherwise too dangerous for animals their size without protective burrows *and* a high-performing sense of vision. Larger goby species that are capable of digging burrows on their own (*Valenciennea sp.*) and small species relying heavily on camouflage (*Gnatholepis sp., Heteroplopomus sp.*) share the sandy habitats with the shrimp-associated gobies.

**Figure 1:**
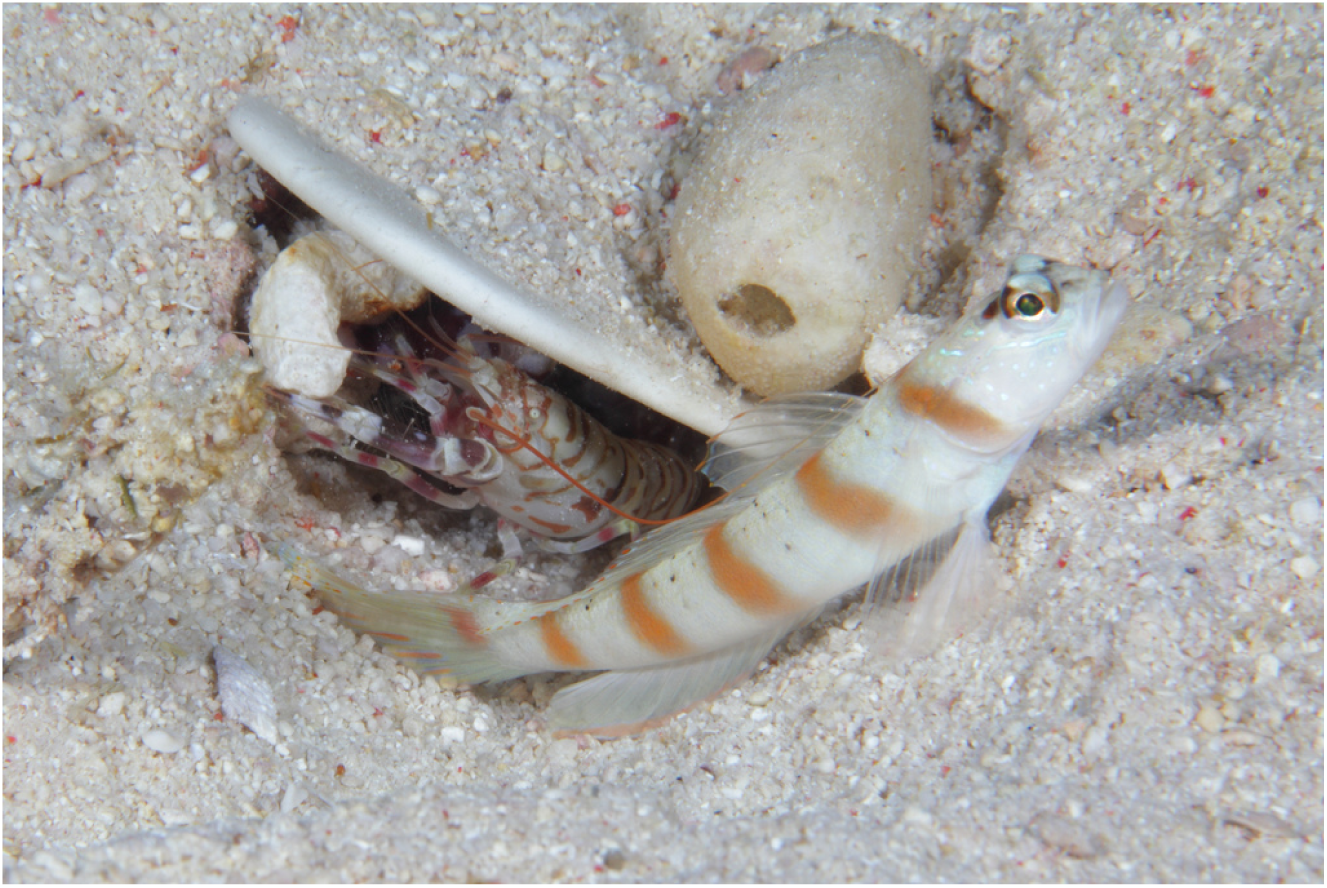
The symbiosis between *Amblyeleotris rubrimarginata* and *Alpheus sp.:* The goby keeps watch for approaching predators, while the shrimp excavates the shared burrow. Note the contact of the shrimp antenna with the dorsal fin of the goby. Malapascua, Philippines. Photograph by K.M.S.

The shrimp-goby symbiosis hence is a trade between muscular – terrain modifying service (shrimp) and visual-sensory service (goby). The investment into the benefit of the symbiotic partner is presumably outweighed by a decreased mortality of both the shrimp and the goby on otherwise shelter-less and dangerous sandy plains, roamed by predatory fishes.

We ask what the economics of biological energy usage of this symbiosis are: Does the goby invest into larger eyes for this symbiosis? The retina is energetically very expensive (Wong-Riley, 2010). For the goby, larger eyes (and hence larger retinae) would incur additional energetic costs both in eye development and maintenance. Selective pressures produce a trade-off between costs and benefits of sensory systems like the visual system (Niven & Laughlin, 2008). While the shrimp-associated gobies evolved from sand-living gobies which are equally vision-centric fishes, the symbiosis could have shifted the optimal point of this trade-off. The eyes’ increased energy consumption would then be outweighed by a decreased mortality due to the advantages conveyed to both partners of the symbiosis.

Alternatively, this symbiosis could be energy-neutral, and simply a result of increased behavioral complexity. Knowledge of the energetics of this symbiosis would certainly aid our understanding of its ecology and evolution. Some of the energetic factors in the shrimp-goby symbiosis can be measured, others can be estimated. Here we attempt to estimate the increased investment in vision by the goby attributable to its visual contribution to the shrimp-goby symbiosis, based on a comparative approach.

## Methods

We compiled a table of fish lengths, eye diameters and maximum depths of occurrence for 8575 marine fishes in FishBase (Froese & Pauly, 2007). The morphometric measurements were made from photographs. The photos used were chosen to be representative of the adult of the species. The measurements of eye sizes from photos should have the same the variability as measurements made on actual specimens, and we used the means. Of the 386 species of gobies measured, we have 5 species with 3 measurements and 55 species with 2 measurements, so we do not have enough species to analyze or test the eye size variability.

Fishbase has been as the basis for a number of interesting analyses, like of the colonization of the deep sea by different fish families (Priede and Froese, 2013), fish length-weight relationships (Froese, 2006) and of the socio-economic & ecological patterns underlying biodiversity decline world-wide (Clausen & York, 2007). We believe that for the analysis of large-scale patterns involving multiple species, Fishbase is a powerful tool.

It is unavoidable that a database of this magnitude contains a small number of mistakes and omissions. Certain types of information about some fishes could be unknown, incompletely determined (the maximum occurrence depth could be underestimated) or incorrectly entered from the literature (in rare cases, we believe). A species coverage of less than 100% will not, however, prohibit a meaningful analysis of patterns across many species. Here, we analyze fish length and eye size. As an example, a goby listed at 3 cm total length could possibly grow to 3.2 cm under ideal conditions. This hypothetical mistake in fish length will not significantly affect the analysis of hundreds of species, as there is no indication that such mistakes would introduce a systematic error into our data. The second parameter we analyze, eye size, was specifically determined for this study from photographs. Here, hypothetically a mis-identification of a fish species could lead to a measurement of the eye size of an incorrect fish. We believe that this case is rare, and the fish in question would very likely be at least from the correct genus, likely with a very similar eye size. In conclusion we believe that while we can’t completely exclude imperfections of our data, they will be rare, small and not systematic.

For this study, we analyzed the gobiidae only. We analyzed 73 shrimp associated species of the genera *Amblyeleotris, Cryptocentrus, Ctenogobiops, Mahidolia, Myersina, Stonogobiops, Tomiyamichthys and Vanderhorstia.*

Similarly, we analyzed 313 solitary species of the genera *Aboma, Acentrogobius, Akko, Amblychaeturichthys, Amblygobius, Amoya, Aphia, Apocryptodon, Arenigobius, Aruma, Asterropteryx, Astrabe, Aulopareia, Austrolethops, Barbulifer, Barbuligobius, Bathygobius, Boleophthalmus, Bollmannia, Bryaninops, Cabillus, Caffrogobius, Callogobius, Chaenogobius, Chriolepis, Chromogobius, Clariger, Clevelandia, Coryogalops, Coryphopterus, Cristatogobius, Cryptocentroides, Ctenogobius, Deltentosteus, Didogobius, Discordipinna, Drombus, Ego, Elacatinus, Eleotrica, Eucyclogobius, Eugnathogobius, Eutaeniichthys, Evermannia, Eviota, Evorthodus, Exyrias, Favonigobius, Feia, Fusigobius, Gillichthys, Gladiogobius, Glossogobius, Gnatholepis, Gobiodon, Gobionellus, Gobiopsis, Gobiosoma, Gobius, Gobiusculus, Gobulus, Gymneleotris, Gymnogobius, Hetereleotris, Istigobius, Koumansetta, Lepidogobius, Lesueurigobius, Lesueurigobius, Lubricogobius, Luciogobius, Lythrypnus, Macrodontogobius, Mauligobius, Microgobius, Millerigobius, Mugilogobius, Odondebuenia, Odontamblyopus, Oligolepis, Ophiogobius, Oplopomus, Opua, Oxyurichthys, Palutrus, Pandaka, Parachaeturichthys, Paragobiodon, Parapocryptes, Paratrypauchen, Parkraemeria, Parrella, Periophthalmus, Platygobiopsis, Pleurosicya, Pomatoschistus, Ponticola, Priolepis, Psammogobius, Pseudogobius, Psilogobius, Pterogobius, Pycnomma, Redigobius, Rhinogobiops, Risor, Sagamia, Scartelaos, Signigobius, Silhouettea, Speleogobius, Sueviota, Sufflogobius, Thorogobius, Tridentiger, Trimma, Trimmatom, Trypauchen, Valenciennea, Yongeichthys, Zebrus and Zosterisessor.*

Some of the fish lengths were given as standard length (SL, anterior tip to the tail base), others as total length (TL). We conducted both an analysis where we excluded TL (the minority of the data points), and where we converted TL to SL by multiplying by 0.8, which is a good estimate of the relationship of TL to SL for gobiidae (based on our measurements of photographs). The results were qualitatively identical, and we only report results of our analysis that included data-points with SL converted to TL.

The size of a fish eye is thought to be a good proxy for the size of the retina (Lee & Stevens, 2007). The retina is energetically expensive (Wong-Riley, 2010), and the retina size is likely what determines the energy cost of the fish visual system. Our analysis is agnostic about visual processing, especially downstream of the retina. We can not make any statements about how the gobies use the information contained in the light their eyes collect. We can however, via the proxy of the retina size, get a good estimate for the energy use of their visual systems.

## Results

We correlated the goby length with the goby eye size for shrimp-associated and solitary gobies (Fig. 2). The regression for the shrimp-associated gobies was 0.031 x + 0.0895, R^2^ = 0.539, and for the solitary gobies 0.0303 x + 0.104, R^2^ = 0.626. The distributions were not significantly different (Mann-Whitney U-Test, p > 0.05). Hence, the eyes of shrimp-associated gobies are not bigger than the eyes of solitary gobies of the same body length. **Symbiosis-caused additional energy usage of the development and operation of the visual system of shrimp gobies is therefore likely small to non-existent.**

**Figure 2:**
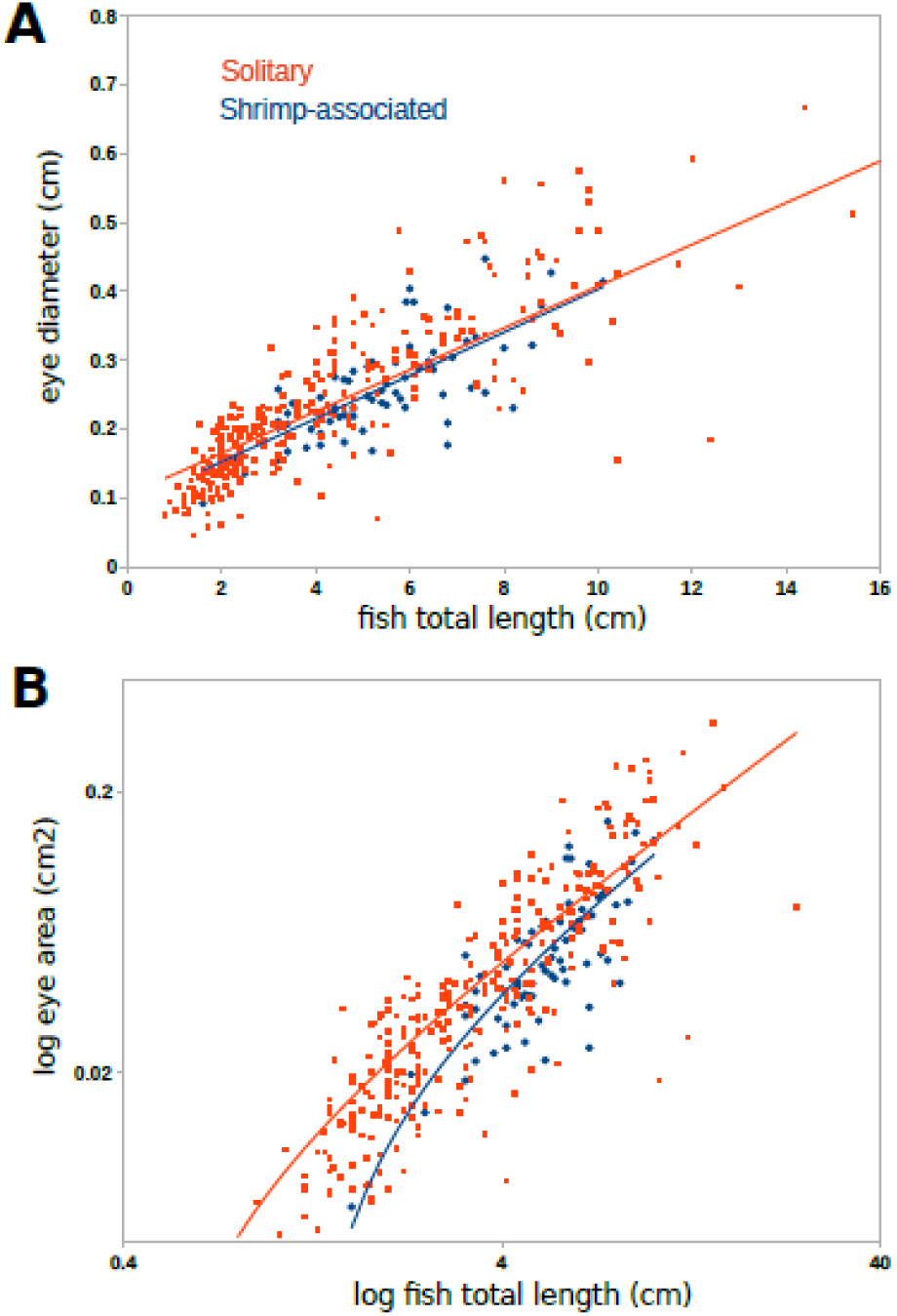
Top: Correlation of fish length with eye diameter for solitary gobies (red squares, red linear regression line) and shrimp-associated gobies (blue diamonds, blue linear regression line). Four of the largest solitary species (length > 10 cm) are excluded on the graph for scaling purposes. Bottom: The same data, but with the retina area calculated from the diameter, plotted against fish length on logarithmic axes, to emphasize comparison over different size classes (Rainer Froese, personal communication, 2016).

To show that the eye diameter/fish length ratio which we here use as a proxy for energy invested into the visual system is meaningful in terms of sensory ecology, we also plotted this ratio for the Muraenidae (moray eels). These fishes are known to be nocturnally hunting olfactory specialists, with little reliance on vision. In order to compensate for the difference in body shapes between Muraenidae and Gobiidae, we calculated the body weight according to the equation:

*weight = a length*^*b*^

For the Gobiidae we chose a=0.0059 and b=3.13 and for the Muraenidae a=0.0011 and b=3.0, according to a Bayesian analysis of fish weight – length relationships (Froese et al. 2014; Froese & Pauly, 2014). This analysis shows that indeed the eye diameter/calculated fish weight ratio is much smaller for the Muraenidae than for gobies (Fig. 3, regression 1.73 10^−3^x + 0.38, R^2^ = 0.162), pointing towards a sensory-ecological relevance of relative eye sizes.

**Figure 3:**
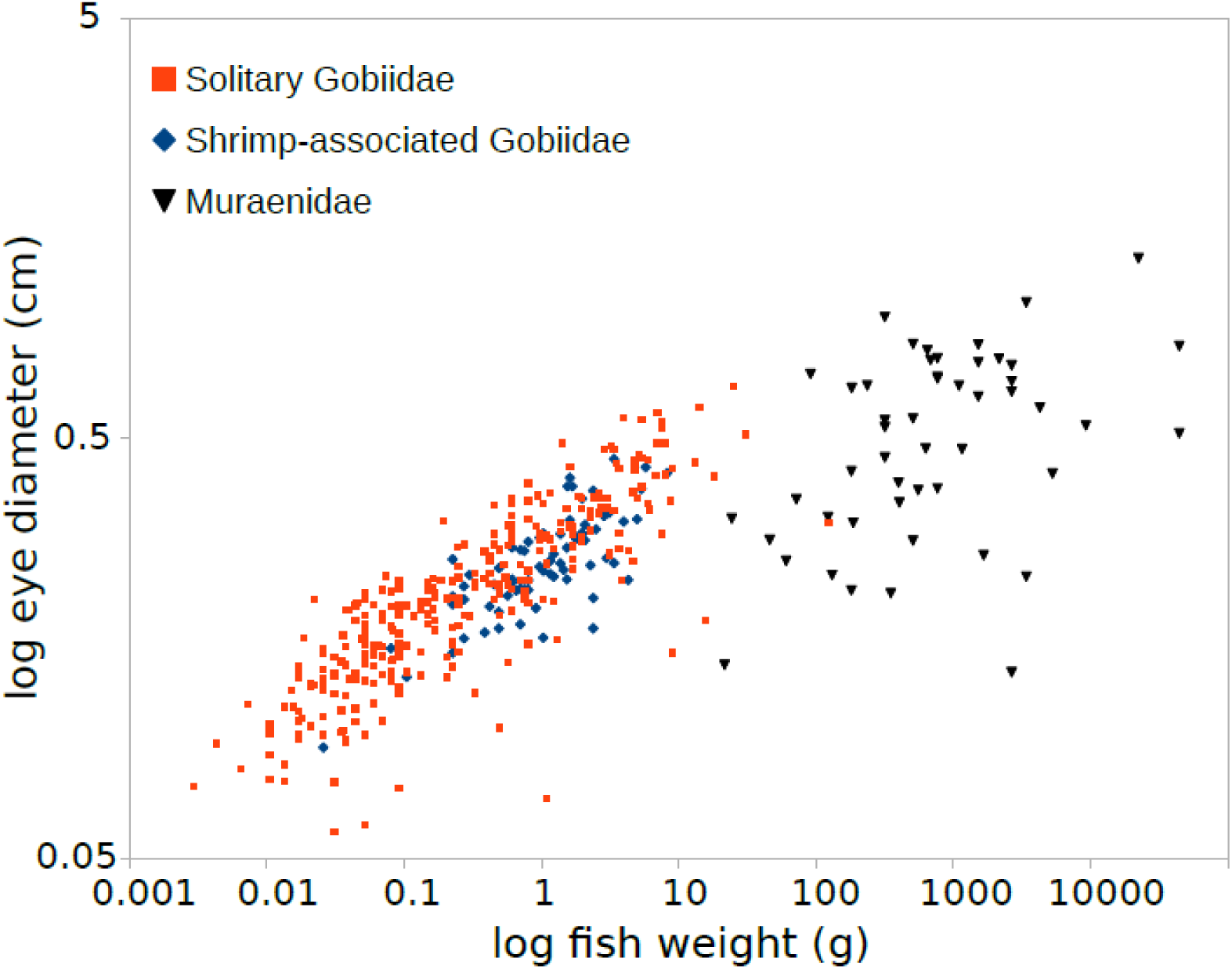
Correlation of calculated fish weight with eye diameter for solitary gobies (red squares) and shrimp-associated gobies (blue diamonds), as well as for the moray eels (*Muraenidae*, black triangles), plotted on logarithmic axes. The small eye diameter in relation to the fish length of the olfactory specialist *Muraenidae* shows the sensory ecological relevance of the eye diameter/fish length relationship. The fish weight was estimated from the fish lengths according to w = a L^b^, with family specific Bayesian-best-fit values for the Gobiidae (a=0.0059 and b=3.13) and the Muraenidae (a=0.0011 and b=3.0; Froese *et al.* 2014; Froese & Pauly, 2015).

We also investigated the depth distribution of the shrimp-associated and the solitary gobies. If the goby-shrimp symbiosis is primarily visual, then it should be confined to shallower depths, at which light levels are significant. We are aware that the depth data for small, often cryptic fishes is less than ideally reliable. Still, some trends emerge. The deepest solitary goby in our data is a pelagic goby, *Sufflogobius bibarbatus*, at 340 m, The deepest shrimp-associated goby is *Stonogobiops xanthorhinica*, at 45 m. The maximum depth/species number distributions of the solitary and symbiotic gobies are also significantly different (p < 0.05, Fig. 4).

**Figure 4:**
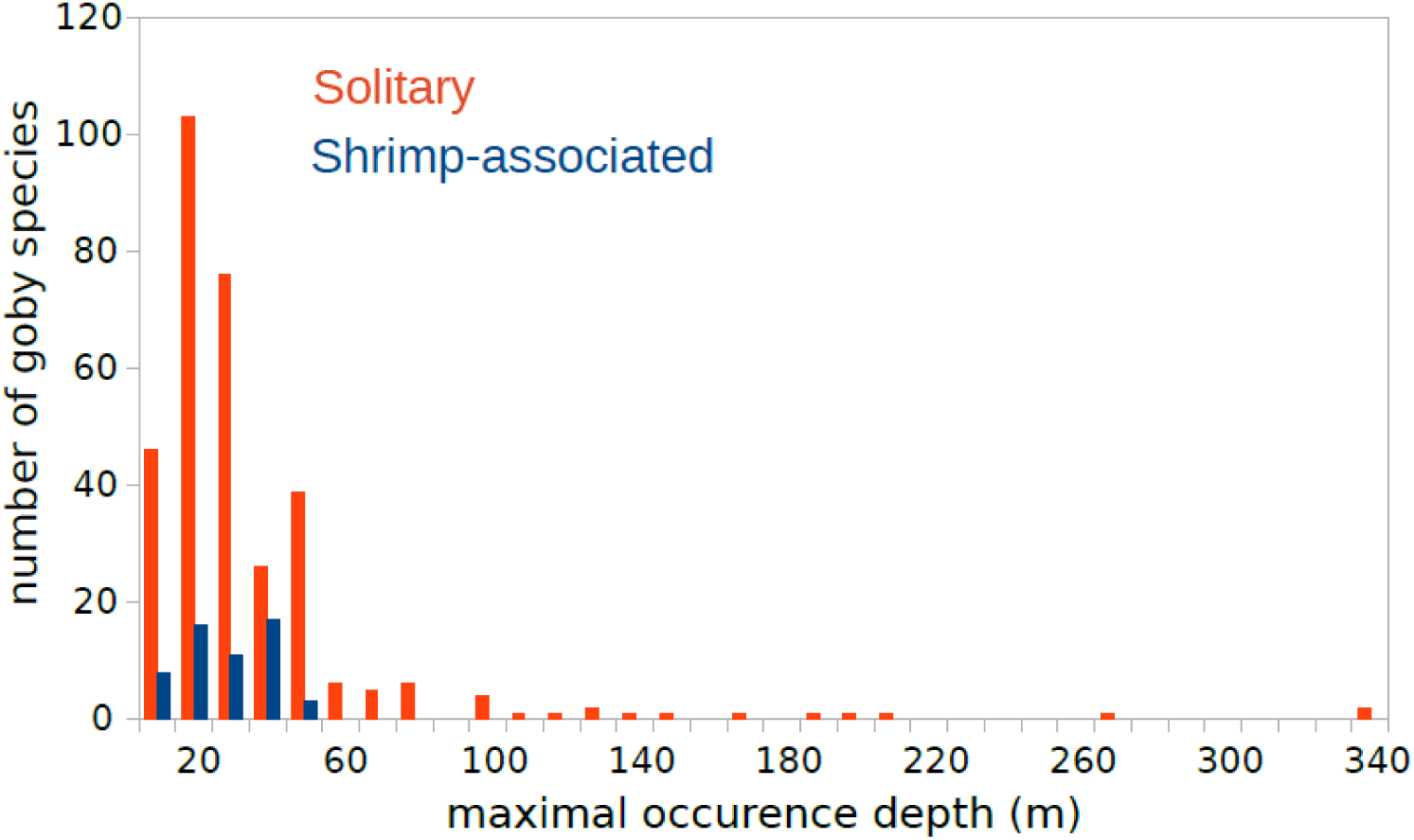
Histogram of the depth distribution of shrimp-associated gobies (blue) and solitary gobies (red). Bin size is 10 depth meters. According to FishBase, the deepest living shrimp-associated goby is *Stonogobiops xanthorhinica* at 45 m, and the deepest living solitary goby is *Sufflogobius bibarbatus* at 340 m.

Sources from outside of our database corroborate this trend: The deepest maximum depth for shrimp-associated gobies listed in Allen et al. (2005) is 48 m for *Amblyeleotris randalli*. We have observed *A. randalli* and *A. guttata* at a depth of 50 m (personal observation, K.M.S, Malapascua, Cebu, Philippines). In a previous study of the goby fauna of Malapascua island, Cebu province, Philippines, the deepest shrimp-associated goby we found was *Amblyeleotris sp.* (“eyebrow”, according to Allen et al., 2005), at 40 m (the survey covered depths to 60 m, Stiefel et al., 2014). In contrast, deep water solitary gobies have been observed to live much deeper, such as *Crystallogobius linearis*, 400 m, *Buenia jeffreysii* (330 m), *Lesuerigobius heterofasciatus*, 345 m and *Gobius roulei*, 385 m (Kovacic & Patzer, 2011). The blind goby *Karsten totoyensis* was collected at a depth of 1122 m and is the deepest goby ever found (Murdy, 2002).

### The preferred occurrence of the symbiosis in shallower (brighter) habitats further supports the fact that vision is a primary factor in the shrimp-goby symbiosis

Alternatively, predation pressure could be lower at depth (Oji, 1996), leading to a diminished need for a burrow and a shrimp-goby symbiosis. A third possibility is that the physical ocean properties (currents) are not favorable for burrow construction at depth; these have been shown to influence the occurrence of gobies (Depczynski & Bellwood, 2005), and especially shrimp-associated gobies (Stiefel et al., 2014). However, this is not likely since in many locations the forces exerted by water (waves, surges, currents) are stronger and more irregular at *shallower* depths, which would cause a distribution of shrimp-associated gobies inverted to the one we observe.

## Discussion

Using a comparative approach, we have demonstrated that the eyes of shrimp-associated gobies are not larger than those of size-matched solitary gobies. Comparing gobies to moray ells, we showed that the fish-size versus eye-size ratio can be ecologically meaningful. We also showed that shrimp-associated gobies are restricted to shallower waters (<50 m). We are aware that senses other than vision (such as audition) contribute to the alarm-behavior of the gobies (for an excellent review of the neural wiring in the escape system of bony fishes see Medan & Preuss, 2014). We do not study these senses here, and it remains a possibility that they are evolutionarily expanded in response to the guard-role of the goby in the symbiosis.

Comparing the regressions of eye size versus fish size is the simplest, and hence most preferable method for the phylogenetic comparison attempted in this study. More complex methods, like the paired contrasts phylogeny method (Maddison, 2000) are not necessary when comparing a feature among a group of closely related species, might actually introduce artifacts due to some of its assumptions (Maddison, 2000) and fail to pick up correlations (Grafen & Ridley, 1996).

Our results show that the shrimp-goby symbiosis likely incurs low or none additional energy costs from the visual system of the goby. Other costs of the symbiosis include reduced foraging due to time spent as a lookout by the goby (see Lyons, 2012). The communication of warnings by the goby to the shrimp alone is most likely energetically cheap. Low energy costs would have several potential evolutionary implications. Presumably, it would make this symbiosis easier to evolve, and indeed it has been shown that it evolved twice in the Indo-Pacific (Thacker et al., 2011). In addition, a goby in the eastern Atlantic is in a symbiotic/commensal relationship with an axiid shrimp (*Axiopsis serratifron*, Wirtz, 2008).

A number of additional factors seemed to be conductive for the emergence of the shrimp-goby symbiosis: 1. Burrow-living in gobies is not unusual. Some of the sister genera of the shrimp-associated gobies (Thacker et al., 2011) hide in crevices (*Callogobius sp.*) or hide in burrows they dig themselves (*Valencienna sp.*). 2. Furthermore, gobies are a very species-rich family. The rapid speciation (Thacker et al., 2011; Rüber et al., 2003) likely helped in the evolution of the shrimp-goby symbiosis, by giving evolution more “shots” at achieving this symbiotic combination. 3. The other partner in the symbiosis, the Alpheid shrimp, generally seem to be conductive to enter mutualistic relationships, inter- and intra-species. Besides the shrimp-goby symbiosis, a symbiosis between an alpheid shrimp and a xanthoid crab in North American salt marshes is also known (Silliman et al., 2003). The sponge-dwelling eusocial Alpheid shrimps (*Synalpheus sp.*) provide an example by this group of a close *intra*-species mutualistic relationship (Macdonald et al., 2006).

## Acknowledgments

Both K.M.S and R.B.R. analyzed the data and wrote the manuscript, R.B.R. provided the database. We thank Dr. Rainer Froese, Dr. Greg Cohen, Dr. Alex Holcombe and Dr. Dario Protti for discussion of the topics covered in this paper. This is FIN contribution number 204.

